# Zinc-independent activation of Toll-like receptor 4 by S100A9

**DOI:** 10.1101/796219

**Authors:** Andrea N. Loes, Ran Shi, Michael J. Harms

## Abstract

The homodimer formed by the protein S100A9 induces inflammation through Toll-like receptor 4 (TLR4), playing critical roles in both healthy and pathological innate immune responses. The molecular mechanism by which S100A9 activates TLR4 remains unknown. Previously, the interaction between purified S100A9 and TLR4 was shown to depend on Zn^2+^; however, the Zn^2+^ binding site(s) on S100A9 were not identified. Here, we investigated the role of Zn^2+^ binding in the pro-inflammatory activity of S100A9. We found that the S100A9 homodimer was prone to reversible, Zn^2+^-dependent aggregation *in vitro*. Using a combination of site-directed mutagenesis and Isothermal Titration Calorimetry (ITC), we identified multiple residues that contribute to Zn^2+^ binding in S100A9. We then used mutagenesis to construct a version of S100A9 with no detectable Zn^2+^ binding by either ITC or Inductively Coupled Plasma-Mass Spectrometry. This protein did not exhibit aggregation upon addition of saturating Zn^2+^. Further, despite the lack of Zn^2+^-binding, this protein was capable of activating TLR4 in a cell-based functional assay. We then modified the functional assay so the Zn^2+^ concentration was exceedingly low relative to the concentration of S100A9 added. Again, S100A9 was able to activate TLR4. This reveals that, despite the ability of S100A9 to bind Zn^2+^, S100A9 does not require Zn^2+^ to activate TLR4. Our work represents an important step in clarifying the nature of the interaction between S100A9 and TLR4.

## INTRODUCTION

The protein S100A9 potently activates inflammation through Toll-like receptor 4 (TLR4) [1]. S100A9 is found at high levels in the extracellular space in inflamed tissues, both as a homodimer and as a heterodimer with the protein S100A8 [2,3]. When properly regulated, S100A9 recruits neutrophils and promotes physiological processes such as angiogenesis [4,5]. Excessive S100A9 can, however, lead to pathological inflammation [1,6]. S100A9 is overexpressed in a variety of inflammatory diseases, where it increases the severity of the inflammatory response [1,7–10]. In extreme cases, it can form a powerful feedback loop with other proinflammatory stimuli, leading to septic shock [6,11]. S100A9, and its close evolutionary relatives S100A8 and S100A12, have therefore been identified as potential therapeutic targets for treating inflammatory disorders [8].

Despite the biological importance of the S100A9/TLR4 interaction, its molecular basis remains poorly understood. Some have proposed that activation occurs via a direct, stoichiometric binding interaction between S100A9 and the TLR4 signaling complex [8]. Others have proposed that higher order oligomers of S100s may activate TLR4[12,13] and that aggregation of S100A9 may play a role in regulation of TLR4 activation[14,15]. Individual mutations to S100A9 have been used to argue in favor of a direct interaction via the canonical target binding interface of S100s[16], but the evidence remains far from conclusive.

One observation informing these models was that, *in vitro*, the interaction between S100A9 and the TLR4/MD-2 complex required both Ca^2+^ and Zn^2+^ ions [8]. S100A9, like other members of the S100 family, has two Ca^2+^ binding sites per monomer. It has multiple residues that may act as ligands for Zn^2+^, but the site of binding is unknown. Transition-metal binding by the heterodimer has been well characterized[17,18], as the heterodimer’s antimicrobial activity is metal-dependent. Several of S100A9 residues do interact with transition metals in the heterodimer formed with S100A8, hinting that they may play a similar role in the homodimer. However, despite a potential role for Zn^2+^ binding in regulation of TLR4 activation by S100A9, little work has been done to identify how homodimeric S100A9 binds to Zn^2+^, nor how Zn^2+^ binding regulates its biological functions.

To gain insight into the mode of interaction between S100A9, TLR4, and Zn^2+^, we set out to identify the sites of Zn^2+^ binding on human S100A9. We identified several sites on the S100A9 homodimer that participate in Zn^2+^ binding. Mutagenesis experiments that remove these key residues completely ablate Zn^2+^ binding and metal-dependent aggregation. Surprisingly, a mutant of S100A9 with no detectable Zn^2+^ binding activates TLR4 in a cell-based assay. This suggests that Zn^2+^ binding is not necessary for the interaction of S100A9 with TLR4.

## RESULTS

### The S100A9 homodimer binds to Zn^2+^ under conditions in which it activates TLR4

We first set out to determine whether S100A9 indeed binds Zn^2+^ under conditions in which it activates TLR4, as the previous observation was limited to an *in vitro* TLR4/MD-2 binding assay [8]. We measured activation by transfecting HEK293T cells with plasmids encoding TLR4 and its cofactors MD-2 and CD14. HEK cells do not natively express TLR4, so activation is strictly dependent on the transfected TLR4/MD-2/CD14 gene products [19]. We also transfected a plasmid encoding a luciferase gene downstream of an NF-kB response element. The total luciferase output of the cells thus corresponds to the degree of TLR4-induced NF-kB activation[19,20].

As a positive control, we added lipopolysaccharide—the canonical ligand of TLR4—and observed a potent activation of NF-kB signaling (Fig 1A). This response could be ameliorated with the addition of polymyxin B (PB), which binds LPS and prevents its binding to the TLR4/MD-2 complex (Fig 1A). We then added recombinantly expressed and purified human S100A9 in the presence of PB. As has been observed previously[1,6,8,21–23], recombinant S100A9 potently activated the NF-kB response, even in the presence of excess polymyxin B (Fig 1A).

**Fig 1.**
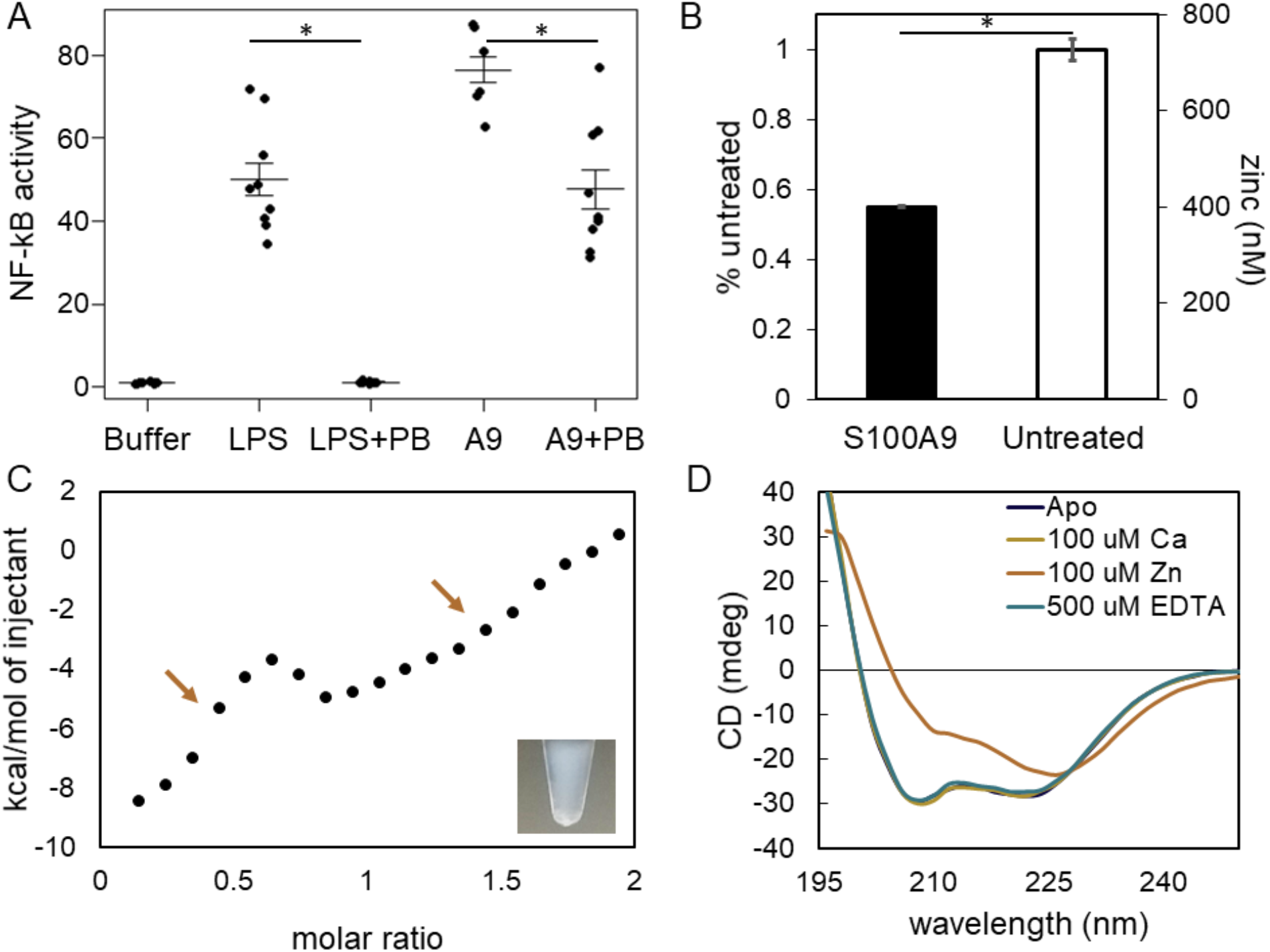
The S100A9 homodimer binds to Zn^2+^ under conditions in which it activates TLR4. A. Activation of TLR4 by S100A9 occurs in the presence and absence of polymixin B. Lipopolysaccharide (LPS) is used as a positive control for expression and activation of the complex. Polymyxin B is included to control for endotoxin-mediated activation of the complex. Points are the technical triplicates from three biological replicates. Horizontal lines are the mean of the biological replicates. Error bars are SEM of the biological replicates. B. S100A9 depletes zinc from cell culture media. The concentration of zinc in cell culture media in S100A9-treated and untreated DMEM + FBS as measured by ICP-MS. Error bars show standard deviation between duplicate samples. Statistically significant differences (single-tailed Student’s t-test) are noted on each panel (* p < 0.05) C. S100A9 binds to zinc. Plot shows integrated heats from isothermal calorimetry experiment. Inset shows protein aggregation occurred during zinc titration. Arrows indicate two phases of zinc binding. D. Zn^2+^ induces reversible aggregation in S100A9. Addition of Zn^2+^ (orange) leads to aggregation of S100A9 and loss of signal for α-helical secondary structure as measured by far UV circular dichroism. Addition of EDTA, restores helical signal (teal).

We next asked whether S100A9 bound to Zn^2+^ under these assay conditions. We added S100A9 to the media that we used to grow HEK cells. We then ran the treated media through a 3K filter—which should remove protein, but not free metal—and measured the zinc content of the eluate using Inductively-Coupled Plasma Mass Spectrometry (ICP-MS). We found that the S100A9-treated sample depleted the zinc concentration by 50% relative to a control that underwent mock treatment without S100A9 (Fig 1B). This suggests that S100A9 does, indeed, bind to Zn^2+^ under these conditions.

### Zn^2+^ binds S100A9 in vitro and induces reversible aggregation

We next set out to study Zn^2+^ binding under more tightly controlled *in vitro* conditions. We followed the binding of Zn^2+^ to S100A9 using Isothermal Titration Calorimetry (ITC) under conditions approximating the extracellular environment (pH 7.4, 100 mM ionic strength, and 200 uM Ca^2+^). The reaction was exothermic and exhibited a shallow change in heat with increasing Zn^2+^ concentration (Fig 1C). The curve exhibits two, visually distinct, phases. The first is a high magnitude exothermic process, followed by a second, lower-magnitude exothermic process (indicated by arrows in Fig 1C). This contrasts sharply with the published ITC results for the interaction between Zn^2+^ and the heterodimer of S100A8/A9, which exhibits an extremely sharp, 2:1 binding transition corresponding to two individual sites with nanomolar binding affinity [18]. These ITC results are difficult to interpret with a thermodynamic binding model, as binding was also accompanied by massive aggregation (Fig 1C, inset). The aggregation behavior of S100A9 in response to Zn^2+^ was extremely robust. We altered buffer conditions, Zn^2+^ counter ions, protein concentration, as well Ca^2+^ concentration. Aggregation occurred within seconds for all conditions at or above equimolar concentrations of Zn^2+^ to S100A9.

Strikingly, however, the Zn^2+^-dependent aggregation of S100A9 was reversible: upon addition of EDTA, the aggregated protein rapidly returned to its soluble form. We can observe the reversibility of this process by eye for the highest concentrations we investigated (1 mM ZnCl_2_, Supp Video). We can also quantify the recovery by far-UV circular dichroism (CD). Protein aggregation is associated with a loss of helical signal upon the addition of Zn^2+^ (Fig. 1D). Upon addition of EDTA, the solution clears and the helical signal returns (Fig 1D). This indicates we are observing near complete recovery of protein secondary structure upon addition of the chelator.

### Disruption of Zn^2+^ binding

These observations are consistent with the previous proposal that the interaction between S100A9 and TLR4 is Zn^2+^-dependent—possibly through an aggregated state. We therefore set out to test the hypothesis that Zn^2+^ binding is necessary for the activation of TLR4 by S100A9. To do so, we set out to identify and disrupt the site(s) of Zn^2+^ binding in S100A9. Using the structures of other S100 proteins, evolutionary conservation, and chemical reasoning, we identified three possible sets of Zn^2+^ ligands that could form (possibly overlapping) binding sites on the S100A9 homodimer (Fig 2).

**Fig 2.**
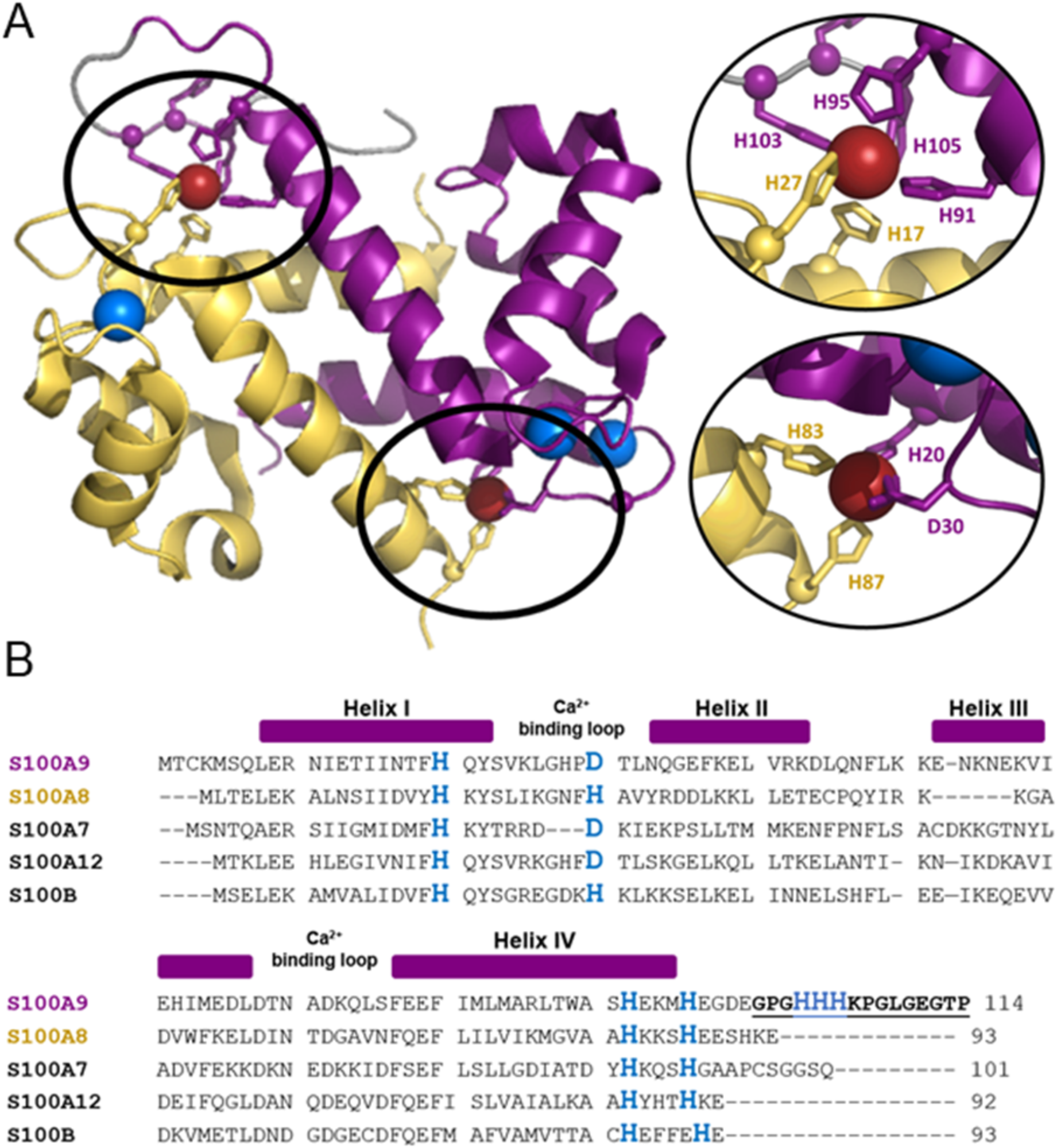
Zn^2+^-binding by heterodimer of S100A8/S100A9 provides hypotheses for metal binding residues in S100A9 homodimer. A. Cartoon representations of crystal structure for S100A8(yellow)/S100A9(purple) heterodimer (PDB:4ggf)[17] in the presence of calcium and manganese. The C-terminal tail (grey) of S100A9 is disordered in the absence of transition metals but becomes structured when histidines in the tail coordinate metal ions. Two distinct interdimer metal binding sites are present in the heterodimer as shown. B. Multiple sequence alignment of S100 proteins reveals conserved residues that may bind Zn^2+^ (blue).

The first potential ligand was the single Cys residue in the protein (Cys3). This Cys is conserved across many S100 proteins. It is part of a short, disordered region of the N-terminus and could plausibly form an interdimer metal Zn^2+^ site, thereby inducing aggregation. Further, this specific Cys is important for metal-dependent aggregation in other S100 proteins [24,25]. We substituted Ser for Cys at this position and remeasured binding by ITC. The C3S protein, however, remained aggregation-prone and was essentially indistinguishable from wildtype human S100A9 (Fig 3A).

**Fig 3.**
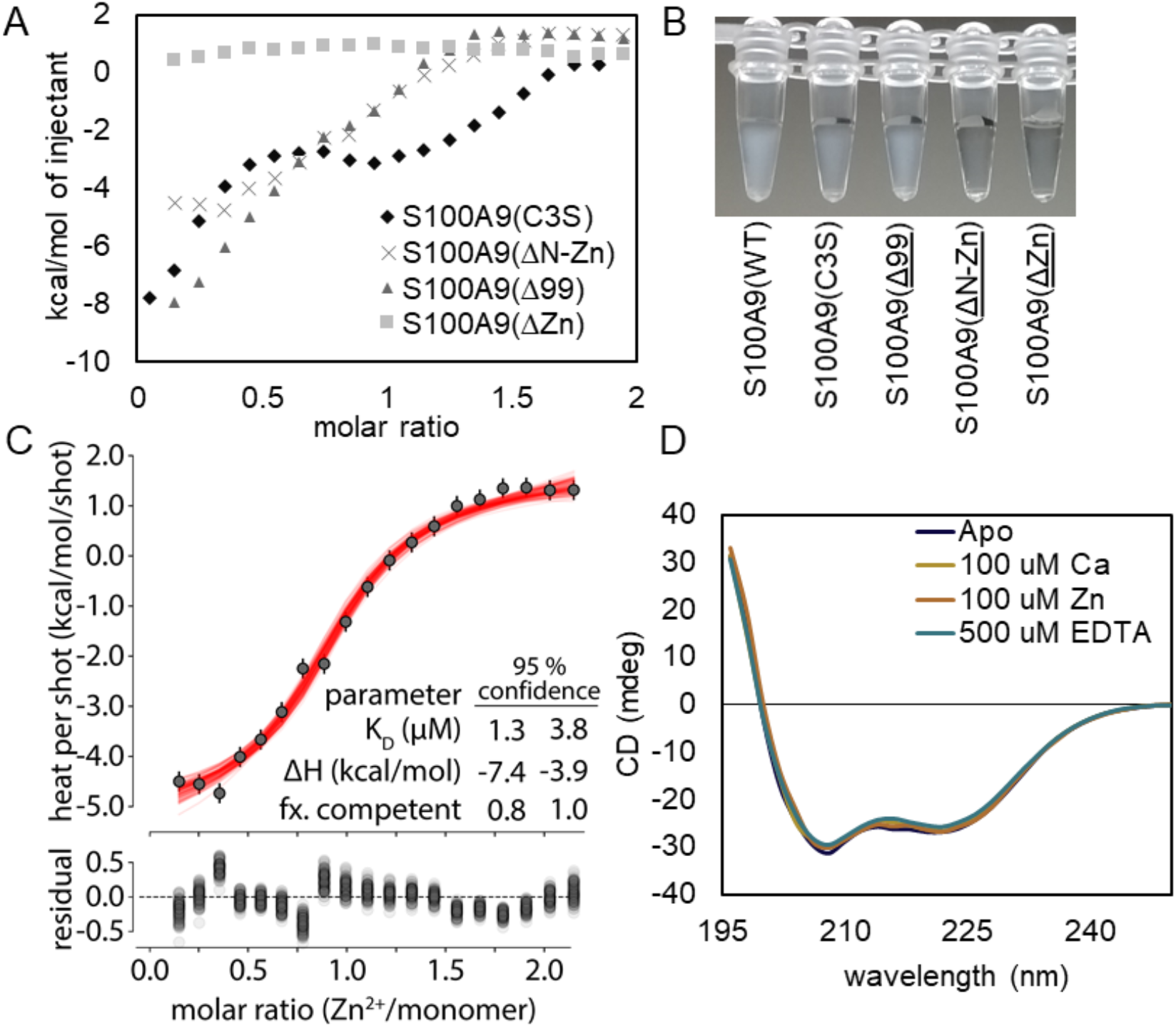
Mutagenesis of S100A9 disrupts Zn^2+^ binding and Zn^2+^-dependent aggregation. A) Points show heats observed as Zn^2+^ was titrated onto S100A9 metal binding mutants: S100A9(C3S) (diamonds), S100A9(Δ99) (triangles) and S100A9(ΔZn) (squares), and S100A9(ΔN-Zn) (crosses). B. Zn^2+^ dependent aggregation of S100A9 is disrupted by mutagenesis of metal binding residues. Photograph shows protein solution post Zn^2+^ titration in ITC. Aggregation did not occur for S100A9(ΔN-Zn) and S100A9(ΔZn). C. A single-site binding model fit to the ΔN-Zn data. Lines show results sampled from 10,000 Monte Carlo samples of possible fits using pytc [26]. Values are lower- and upper-bounds for 95% credibility interval. D. Addition of Zn^2+^ to S100A9(ΔZn) does not result in changes to α-helical secondary structure as measured by far UV circular dichroism.

Another potential collection of Zn^2+^-binding ligands are part of a C-terminal extension unique to S100A9. This “tail” is disordered in crystal structures of the S100A9 homodimer but is ordered in structures of the S100A8/S100A9 heterodimer bound to transition metals. The tail has three histidines that could conceivably participate in transition metal binding by the S100A9 homodimer (Fig. 2B)[17,18,27]. To test for the importance of these residues for Zn^2+^-binding, we truncated the C-terminal tail of the protein between residues 99 and 100 in the C3S background S100A9(Δ99). This removes three histidine residues, two of which participate in coordination of metals in the S100A8/A9 heterodimer. This mutation did alter the Zn^2+^ binding isotherm measured by ITC (Fig 3A). The titration lost its second phase and, superficially at least, resembles single-site binding. As with the wildtype protein, however, this protein aggregated significantly (Fig 3B).

The final site we attempted to disrupt consisted of residues that form the canonical tetrahedral transition metal site observed in many S100 protein family members. If the S100A9 site is similar to other S100 proteins, it would consist of residues from the N-terminus of one monomer (H20 and D30), and the C-terminus of the other monomer (H91 and H95). To disrupt this putative site, we introduced two mutations into the C3S background: H20N/D30S. We then tested metal binding for this “ΔN-Zn” mutant. Like the truncation mutant, these mutations disrupted of one of the processes involved in Zn^2+^-binding by S100A9 (Fig. 3A). This suggests that the two processes for Zn^2+^ binding observed in the wild-type protein might be occurring separately: one via the predicted tetrahedral Zn^2+^-binding site and the other via the C-terminal extension. Interestingly, we did not observe aggregation of the ΔN-Zn mutant in the sample cell (Fig. 3B). This suggests that the Zn^2+^-dependent aggregation of S100A9 is mediated by Zn^2+^ binding to the canonical Zn^2+^ binding site of S100 proteins. Because of the lack of aggregation, we were able to extract thermodynamic parameters for this experiment (Fig. 3C). The data fit a single-site binding model, suggesting that S100A9 ΔN-Zn binds with a 1 Zn^2+^:1 monomer stoichiometry. The 95% credibility interval for the K_D_ was 1.3-3.8 μM, while the buffer-dependent enthalpy was estimated to be between −7.4 and −3.9 kcal/mol (Fig 3C).

As evidenced by the altered ITC results, both the C-terminal tail and H20/D30 contribute to binding, but removal of either is insufficient to completely disrupt binding. Because our goal is to completely disrupt Zn^2+^ binding to test for its necessity in TLR4 activation, we thus constructed an eight-way mutant protein in which we disrupted all of the residues that may ligate transition metals in S100A9: *C3S/H20N/D30S/H91N/H95N/H103N/H104N/H105N*. We measured Zn^2+^ binding to this eight-way mutant by ITC but found no evidence of binding (Fig 3A). Additionally, S100A9(ΔZn) does not aggregate in the sample cell (Fig. 3B). To verify that adding eight mutations did not disrupt the overall fold of the protein, we examined the far-UV CD signal for this protein and found that it is indistinguishable from that of S100A9, indicating similar secondary structures are present in both proteins (Fig 3D).

### Protein activates without Zn^2+^ binding

Having constructed S100A9 variants with compromised Zn^2+^ binding, we could now test the hypothesis that Zn^2+^-binding was necessary for TLR4 activation by S100A9. We first tested our truncation mutant of S100A9, which lacks the histidine-rich C-terminal tail and appears to exhibit a single-site binding (Fig 3B). We found that S100A9Δ99 activated similarly to wildtype (Fig 4A). Next, we applied our mutant with fully disrupted Zn^2+^ binding to our HEK assay. S100A9(ΔZn) activated TLR4, even without being unable to bind to Zn^2+^ (Fig 4A). To verify that the protein did not bind Zn^2+^ in the experimental conditions, we repeated out attempt to pull Zn^2+^ out of our assay media using S100A9. Unlike the wildtype protein and S100A9Δ99, S100A9(ΔZn) did not pull Zn^2+^ out of the media (Fig 4B).

**Fig 4.**
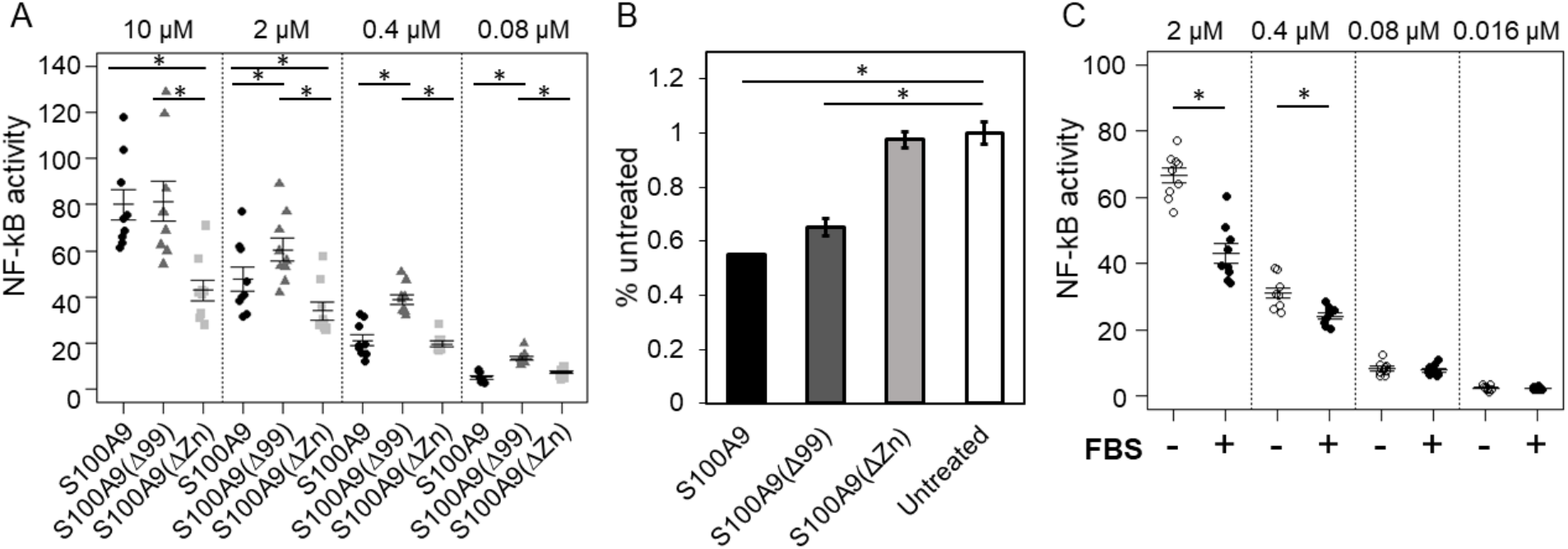
Activation of TLR4 by S100A9 occurs in the absence of Zn^2+^ binding. A. Metal binding mutants of S100A9 activate TLR4. Points are the technical triplicates from three biological replicates. Horizontal lines are the mean of the biological replicates. Error bars are SEM of the biological replicates. B. S100A9(ΔZn) does not deplete Zn^2+^ from cell culture media. The concentration of zinc in cell culture media treated with wild-type S100A9, S100A9(Δ99), and S100A9(ΔZn) as measured by ICP-MS. C. Activation of TLR4 by S100A9 occurs both in high and low Zn^2+^ conditions. Activation by different concentrations of S100A9 in the presence and absence of 10% FBS which contributes Zn^2+^ to the media. Points are technical triplicates from three biological replicates. Error bars show a standard error on the mean. Mean is shown as a horizontal line. Statistically significant differences (single-tailed Student’s t-test) are noted on each panel. * p < 0.05

These results suggest that Zn^2+^ is not necessary for activation. To verify that this holds for the wildtype protein, we re-tested wildtype human S100A9 activity in our TLR4 functional assay without adding Fetal Bovine Serum (FBS). FBS is a typical additive in these assays that brings in several micromolar Zn^2+^ [28,29]. The defined medium without FBS, however, has low concentrations of Zn^2+^ (5 nM)[30]. If Zn^2+^ is not necessary for activation, S100A9 may still activate TLR4 even in the absence of FBS. As predicted, we observed that S100A9 activated to an equivalent level in low-Zn^2+^ FBS(-) media relative to the high-Zn^2+^ FBS(+) media (Fig 4). Indeed, at the highest protein concentrations, S100A9 activated TLR4 slightly more effectively in the FBS(-) vs. FBS(+) sample (p = 0.018).

## DISCUSSION

Here we show that Zn^2+^-binding by the S100A9 homodimer is distinctly different from that of the S100A8/A9 heterodimer and other S100 homodimers. Zn^2+^-binding involves many residues in S100A9, and likely occurs at two sites. One site appears to involve the C-terminal tail and the other the tetrahedral binding site. Zn^2+^-binding also induces reversible-aggregation in S100A9. Intriguingly, we find that Zn^2+^-dependent aggregation of S100A9 is driven by Zn^2+^ binding to the canonical interdimer site on S100A9, not by the disordered C-terminal extension, unique to S100A9.

Retention of activation of TLR4 by the full metal binding site deletion mutant was surprising given previous data suggests that Zn^2+^ is required for S100A9 to interact with the TLR4/MD-2 as purified proteins [8]. However, our results here demonstrate that S100A9 can activate TLR4 independent of Zn^2+^ binding, either as a protein that cannot bind Zn^2+^ (Fig 4A) or in media that does not contain appreciable quantities of Zn^2+^.

The ability to disrupt Zn^2+^ binding without disrupting pro-inflammatory activity may prove extremely useful for dissecting the biology of S100A9. The protein plays to distinct, but related, roles in innate immunity. As a homodimer, it activates inflammation via TLR4[8]. As a heterocomplex with the protein S100A8, it uses high-affinity metal chelation to achieve potent antimicrobial activity [17,18,27,31]. The individual contributions of each function can, however, be difficult to disentangle. For example, to what extent does S100A9 control bacterial growth directly (by nutrient sequestration as a heterocomplex) or indirectly (by activating inflammation and recruiting neutrophils as homodimer)? Until now, such a question would be difficult to answer. Our S100A9(ΔZn) construct could, however, be a powerful reagent to perform such a study, as it remains pro-inflammatory, but also contains multiple mutations known to compromise antimicrobial activity in the heterocomplex[17].

We also show that the C-terminal tail of S100A9 is not required for the interaction with TLR4. Many biological functions both within cells and outside cells have been attributed to the flexible C-terminal tail of S100A9[32–35], the lack of involvement of the C-terminal in the direct interaction with TLR4 suggests that these functions, like antimicrobial activity, may be functionally separable from TLR4-activation by S100A9.

Our results also suggest that the Zn^2+^-dependent aggregation is not required for activation by S100A9. Concentrations of Zn^2+^ in the extracellular space are sufficient to induce aggregation of S100A9 *in vitro*, and S100A9 aggregates have observed in hosts [14,15]. One possibility is that the solution form of S100A9 is active, while the aggregated form is not. If so, aggregation could act as a feedback loop to decrease the pro-inflammatory activity of S100A9. Further work is needed to address whether the Zn^2+^-induced aggregation results in a loss in pro-inflammatory activity.

Finally, Zn^2+^-dependent aggregation of S100A9 has posed a technical challenge to *in vitro* analyses of target binding interactions by S100A9. The observation that activation of TLR4 by S100A9 can occur in the absence of Zn^2+^-binding facilitates further, previously impossible, *in vitro* studies of this.

Our work represents an important step in clarifying the nature of the interaction between S100A9 and TLR4. We have revealed that two structural features of S100A9 appear not to contribute to the S100A9-induced activation of TLR4: Zn^2+^ binding and the disordered C-terminal terminus. Because removal of Zn^2+^ binding also removes metal-dependent aggregation, this also causes us to favor a model in which the soluble form of S100A9—rather than the aggregate—is the species that activates TLR4. Overall, our S100A9(ΔZn) construct is a much simpler molecule than the wildtype form, setting up future studies to identify the structural and mechanistic basis for the activation of TLR4 by S100A9.

## MATERIALS & METHODS

### Plasmids and recombinant protein preparation and characterization

Mammalian expression vectors were obtained from Addgene. The construct containing human TLR4 was made available by Ruslan Medzhitov (Addgene plasmid #13085). Constructs containing human CD14 and ELAM-Luc were obtained from Doug Golenbock (Addgene plasmid #13645 and #13029). Human MD-2 was obtained from the DNASU Repository (HsCD00439889).

A synthetic gene construct for human S100A9 (UniProt #P06702) was designed to be free of common restriction sites and codon optimized and purchased as PUC57 construct from GenScript (New Jersey, USA). S100A9 were cloned into a modified 6xHis MBP LIC TEV vector with NcoI and HindIII to yield a protein constructs with a TEV-cleavable histidine tag. Mutations to S100A9 were made using the QuikChange Lightning Mutagenesis Kit from Agilent Technologies (Santa Clara, CA).

To express S100A9 wild-type and mutants, *E. coli* BL21(DE3) pLysS competent cells containing the expression vectors for S100 proteins were grown overnight at 37°C, diluted 1:150 into LB containing ampicillin and chloramphenicol, grown to mid-log phase (OD_600_ ~ 0.6-1), and induced with 1 mM isopropyl-β-d-1-thiogalactopyranoside overnight at 16° C with aeration. Bacterial pellets were harvested by centrifugation at 3000 rpm at 4° C for 20 min and stored at - 20° C. Pellets (~5 g) were suspended by vortexing and lysed in 25 mL Buffer (25 mM Tris, 100 mM NaCl, 25 mM imidazole, pH 7.4) with 37.5 U DNase I (ThermoFisher Scientific) and 0.75 mg Lysozyme (ThermoFisher Scientific) by shaking at RT for approximately 1 hr. The lysate was clarified by centrifugation at 15,000 RPM for 50 min. at 4° C. Protein was purified using a 5 mL Ni^2+^-NTA HisTrap column from (Healthcare GE) using an FPLC (Akta Biosciences) with gradient elution to HisB (25 mM Tris, 100 mM NaCl, pH 7.4, with 500 mM imidazole for human, mouse and chicken proteins, 1 M imidazole for opossum S100A9). Pooled elution peak of purified protein was cleaved with tobacco etch protease (TEV) overnight at RT. Cleaved protein was collected from a gradient elution of a 5 mL Ni-NTA column from HisA to HisB. Protein purity was assessed with SDS-PAGE and pure fractions were pooled and dialyzed into 20 mM HEPES, 100 mM NaCl, pH 7.4, treated with 10g/L chelex. For proteins containing cysteine (wild-type and S100A9Δ99, 0.5 mM TCEP was included in dialysis. Protein was flash frozen in liquid nitrogen and fresh aliquots were thawed and exchanged into fresh 20 mM HEPES, 100 mM NaCl, pH 7 (prepared with endotoxin-free water) with 3K Nanosep centrifugal devices prior to functional assays. While results shown here are replicates from a single protein prep, TLR4 activation was tested with two independent preps of each mutant to ensure that the results were not batch specific. Protein concentrations were measured using a Bradford assay.

For metal binding experiments, protein was exchanged into 20 mM HEPES, 100 mM NaCl, 500 uM TCEP, 200 uM CaCl_2_, pH 7.4 and concentration was measured with a Bradford assay. Titrations were conducted at 25°C with stirring at 1000 rpm. The titrations were performed all on the same day, with the same titrant (500 uM ZnCl_2_, 20 mM HEPES, 100 mM NaCl, 500 uM TCEP, 200 uM CaCl_2_, pH 7.4) in an ITC200 from MicroCal. Injections were 2 uL. Integrations were performed with MicroCal Software. We extracted thermodynamic parameters for Zn^2+^ binding to ΔN-Zn by fitting a single-site binding model as implemented in pytc[26]. We used the last four points of the titration to account for the heat of dilution. We then generated 900,000 Monte Carlo samples sampling over possible fit values. We reported the 95% credibility interval for the thermodynamic parameters. Note that the “fraction competent” term is a nuisance parameter that captures errors in the concentrations of the titrant or stationary material, as well as the fact that not all protein and/or Zn^2+^ may have been competent to bind. See Freire et al[36].

For secondary structure measurements, protein samples were prepared at 20 μM in 10 mM Trizma, pH 7. Far-UV circular dichroism data was collected between 200–250 nm using a J-815 CD spectrometer (Jasco) with a 1 mm quartz spectrophotometer cell (Starna Cells, Inc. Catalog No. 1-Q-1). Assays for Zn^2+^-dependent aggregation were performed by adding 100 μM CaCl_2_, then 100 μM ZnCl_2_, and finally 500 μM EDTA. Aggregation was tested in duplicate for each protein. Raw ellipticity was converted into mean molar ellipticity using concentration and the number of residues for each protein.

### Cell culture and transfection conditions

Human embryonic kidney cells (HEK293T/17, ATCC CRL-11268) were maintained up to 30 passages in DMEM supplemented with 10% FBS at 37° C with 5% CO_2_. For each transfection, a confluent 100 mm plate of HEK293T/17 cells was treated at room temperature with 0.25% Trypsin-EDTA in HBSS and resuspended with an addition of DMEM + 10% FBS. This was diluted 4-fold into fresh medium and 135 μL aliquots of resuspended cells were transferred to a 96-well cell culture treated plate. Transfection mixes were made with 10 ng of TLR4, 1 ng of CD14, 0.5 ng of MD-2, 0.1 ng of Renilla, 20 ng of ELAM-Luc, and 68.4 ng pcDNA3.1(+) per well for a total of 100 ng of DNA, diluted in OptiMEM to a volume of 10 μL/well. To the DNA mix, 0.5 μL per well of PLUS reagent was added followed by a brief vortex and RT incubation for 10 min. Lipofectamine was diluted 0.5 μL into 9.5 μL OptiMEM per well. This was added to the DNA + PLUS mix, vortexed briefly and incubated at RT for 15 min. The transfection mix was diluted to 65 μL/well in OptiMEM and aliquoted onto a plate. Cells were incubated with transfection mix overnight (18-22 hrs) and then treated with protein (0-10 μM) or LPS (100 ng/well) mixtures prepared in 25% buffer (20 mM HEPES, 100 mM NaCl, pH 7) and 75% cell culture media (DMEM + 10% FBS). *E. coli* K-12 LPS (tlrl-eklps, Invivogen) was dissolved at 5 mg/mL in endotoxin free water, aliquots were stored at −20° C. To avoid freeze-thaw cycles, working stocks of LPS were prepared at 10 ug/mL and stored at 4° C. There has been some concern in testing recombinant DAMPs against TLR4 due to the potential presence of contaminating LPS in proteins which have been expressed in bacteria [37,38]. We tested our S100s in the presence of 50 μg/mL polymyxin B, an LPS binding agent to limit signaling from LPS contamination in recombinant protein preparations. This concentration of polymyxin B eliminated signaling by 100 ng/mL of LPS but had a minimal effect on the signaling by S100A9. Cells were incubated with treatments for 4 hr. The Dual-Glo Luciferase Assay System (Promega) was used to assay Firefly and Renilla luciferase activity of individual wells. Each NF-κB induction value shown represents the Firefly luciferase activity/Renilla luciferase activity.

### Metal depletion experiments

Samples were prepared in the same way as HEK cell treatments. First, 150 uL of 8 uM protein was prepared in chelex treated 20 mM HEPES, 100 mM, pH 7.4, then protein was diluted 1:4 into DMEM + 10% FBS. Samples were incubated in cell culture incubator (37 C with 5% CO_2_) for 1 hr and then filtered with 15 mL 3K microsep columns by centrifuging at 5000 xg for 20 min. Flow-through was collected and sent for analysis by the Elemental Analysis Core at Oregon Health and Sciences University. For each sample 100 μl of the filtered media solution was added to 1 ml of 1 % HNO3 (trace metal grade, Fisher) in a 15 ml acid-rinsed centrifuge tube (VWR). Inductively coupled plasma mass spectroscopy (ICP MS) analysis was performed using an Agilent 7700x equipped with an ASX 500 autos ampler. The system was operated at a radio frequency power of 1550 W, an argon plasma gas flow rate of 15 L/min, Ar carrier gas flow rate of 0.9 L/min. Elements were measured in kinetic energy discrimination (KED) mode using He gas (4.3 ml/min). Data were quantified using a 11-point (0, 0.5, 1, 2, 5, 10, 20, 50, 100, 500, 1000 ppb (ng/g) for Ca, Mn, Fe, Cu, and Zn using a multi-element standard. For each sample, data were acquired in triplicates and averaged. A coefficient of variance (CoV) was determined from frequent measurements of a sample containing ~10 ppb of Mn, Fe, Cu, and Zn. An internal standard (Sc, Ge, Bi) continuously introduced with the sample was used to correct for detector fluctuations and to monitor plasma stability. Elemental recovery was evaluated by measuring NIST reference material (water, SRM 1643f) and found to within 90 - 100% for all determined elements.

## ACKNOWLEDGEMENTS

S1 Movie was kindly shared by Dr. Jeremy Anderson.

## AUTHOR CONTRIBUTIONS

Project was conceptualized by ANL and MJH. Investigation was performed by ANL and RS. Funding acquisition, project administration, resources and supervision were performed by MJH. Visualization and writing of original draft were performed by ANL and MJH. Reviewing and editing were performed by ANL, RS, MJH.

## Supporting information captions

**S1 Movie. Addition of ZnSO_4_ leads to reversible aggregation of S100A9**. The solution shown contains 1 mM S100A9 in 10 mM Tris, 200 mM NaCl, 10 mM CaCl_2_, pH 7.4 at room temperature (approximately 25 °C). Five seconds into the movie, 1 mM ZnSO_4_ is added, leading to massive aggregation. 12 seconds in, 2 mM EDTA is added, dissolving the aggregate.

